# bioLQM: a java library for the manipulation and conversion of Logical Qualitative Models of biological networks

**DOI:** 10.1101/287011

**Authors:** Aurélien Naldi

## Abstract

Here we introduce bioLQM, a new Java software toolkit for the conversion, modification, and analysis of Logical Qualitative Models of biological regulatory networks, aiming to foster the development of novel complementary tools by providing core modelling operations. Based on the definition of multi-valued logical models, it implements import and export facilities, notably for the recent SBML-qual exchange format, as well as for formats used by several popular tools, facilitating the design of workflows combining these tools. Model modifications enable the definition of various perturbations, as well as model reduction, easing the analysis of large models. Another modification enables the study of multi-valued models with tools limited to the Boolean case. Finally, bioLQM provides a framework for the development of novel analysis tools. The current version implements the usual updating modes for model simulation (notably synchronous, asynchronous, and random asynchronous), as well as some static analysis features for the identification of attractors. The bioLQM software can be integrated into analysis workflows through command line and scripting interfaces. As a Java library, it further provides core data structures to the GINsim and EpiLog interactive tools, which supply graphical interfaces and additional analysis methods for cellular and multi-cellular qualitative models.

## 1 Introduction

Logical models are highly abstract dynamical models, which have been proposed to study biological regulatory systems in the late 60s [16, 37]. This modelling framework has since gained popularity [5, 31, 34] and has been successfully applied to a wide range of regulatory and signalling systems [1, 14, 23, 33].

In logical models, components are represented by discrete variables with a small range of possible values, representing qualitative differences in activity. Boolean components can only be active (1) or inactive (0), while multi-valued components define increasing integer activity levels. Regulatory effects are often represented as signed arcs between components in the **regulatory graph**. These effects are further formalised as logical rules (also called logical parameters or logical functions), specifying the target activity level of each component according to the current levels of its regulators (a subset of all model components). While interactive software for model definition such as GINsim [25] or The Cell Collective [15] enable the definition of regulatory graphs, bioLQM solely relies on logical rules. However, signed regulatory arcs can be extracted from these logical rules if needed. The relative simplicity of this formalism enables the definition of large models with dozens of components, without requiring precise knowledge of kinetic parameters. A formal definition of logical qualitative models is provided in appendix A.

The CoLoMoTo consortium was recently founded to facilitate model sharing and foster cooperation in the qualitative modelling community, building on the introduction of the SBML qual exchange format [7, 22]. The bioLQM toolkit presented here reinforces this effort by implementing a collection of model modification and format conversion operations in an extensible architecture illustrated in Figure 1. It can then be used as a building block for novel software development or as a bridge between complementary softwares in complex analysis workflows. It is currently embed in the GINsim software [25], which provides a graphical interface to most of its features. BioLQM is also used as backend for model definition and computation of successor states in Epilog [38], as well as in the CoLoMoTo notebook for model conversion and some dynamical analysis features [26].

**Figure 1:**
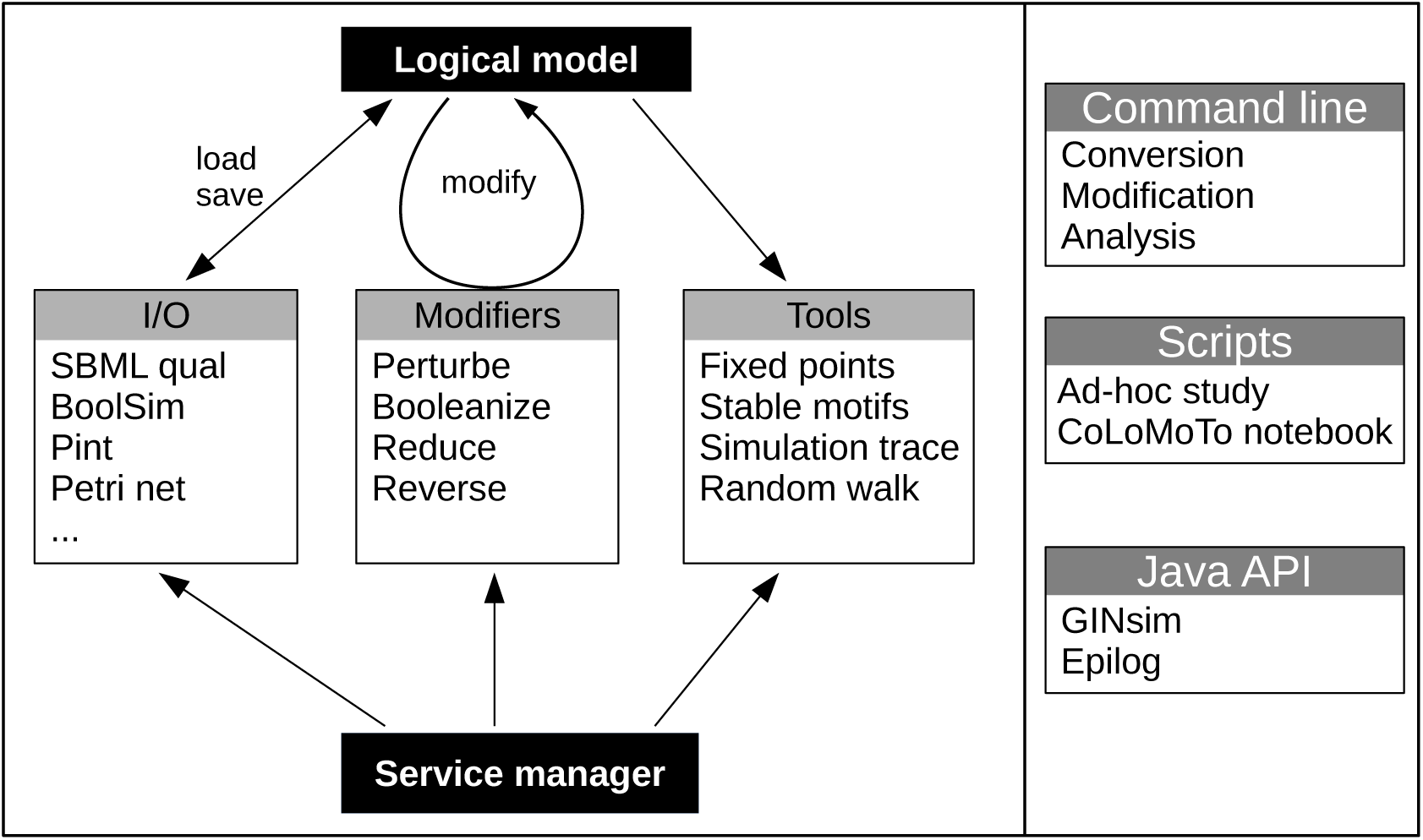
Global structure of the bioLQM toolkit. The bioLQM toolkit is centered around a data structure for the representation of logical qualitative models. Based on this data structure, (i) the `I/O` module contains a collection of formats enabling model loading and saving; (ii) the `modifiers` module contains a collection of model modifiers to transform an input model into a modified model; (iii) the `tools` module contains a collection of analysis tools. All these feature are accessible through a central `service manager`, which handles service discovery and serves as main entry point for the Java API. A simple `command line launcher` provides quick execution of simple workflows, while a `scripting engine` can be used for more complex use cases.

Section 2 introduces model loading, saving and converting operations. Section 3 introduces the simulation and dynamical analysis features. Section 4 introduces model modifications. These features can be accessed from a command-line interface or through a scripting API, as discussed in Section 5.

## 2 Loading and converting Logical Qualitative models

The increasing use of qualitative models to study biological systems led to the development of various software tools for the logical formalism [2, 12, 19, 25, 36] and related qualitative approaches [3, 28, 35]. Many software tools use their own file format for the definition of models, hindering the delineation of analysis workflows combining complementary software tools. The SBML qual exchange format [7] has recently been proposed to improve interoperability between modeling tools. However SBML support is often missing from existing software and is may not be a priority for newer ones.

To ease model exchange between software tools that do not all support the SBML qual format, the bioLQM toolkit provides an extensible list of format handlers connected to the internal model representation. Each format is described as a Java class providing annotations (name of the format, default file extension and multi-valued support) along with optional implementations of model import (loading a file into the internal representation) or export (saving the internal representation to a file) operations. A service manager maintains a list of these descriptor classes through service discovery to facilitate the addition of new formats.

The supported formats are listed in Table 1 and in bioLQM documentation ^1^. BioLQM uses JSBML [30] to load and save SBML qual models. The other import parsers are based on the antlr parser generator [27]. While some formats natively support multi-valued models, many are limited to the Boolean case. Multi-valued models can be exported to these Boolean formats through an implicit booleanization step, described in section 4.

**Table 1:** Available formats. The Import/Export capabilities are listed in the corresponding columns (all formats can be exported). The formats natively supporting multi-valued models are also identified, other formats rely on implicit model booleanization.

| File extension | multi-valued | Import | Export | Description and associated tools |
| --- | --- | --- | --- | --- |
| sbml | x | x | x | SBML qual Exchange format [7] |
| bnet |  | x | x | BoolNet, PyBoolNet [18, 19] |
| booleannet |  | x | x | booleannet [2] |
| boolfunction | x |  | x | Boolean functions |
| boolsim | x |  | x | boolsim (genYsis) [12] |
| cnet | x | x | x | BNS [10] |
| ginml | x |  | x | GINsim [25] |
| tt | x | x | x | Truth table |
| an | x |  | x | Pint automata network [28] |
| apnn, pnml, ina | x |  | x | Conversion to Petri Net formats [6] |
| gna | x |  | x | GNA (Piecewise-linear formalism) [3] |
| bnd |  |  | x | MaBoSS (Stochastic Boolean model) [35] |

## 3 Model dynamics and simulation

A **state** of the model is a vector giving the activity levels of all its components. As the activity level of each component is restricted to a finite range, the **state space** (containing all possible states) itself is also finite. However, the total number of possible states grows exponentially with the number of components. We say that a component is **called to update** in a given state if the evaluation of the associated logical rule is different than its current activity level: for example an inactive component can become active. When analysing logical models, we are often interested in **stable states** (also called fixed points, or steady states), in which no component is called to update. Such stable states denote a qualitative equilibrium in which all components can maintain their current activity level.

The dynamics of the model (*i.e.* its evolution over time) is given by transitions between states states of the model, controlled by the updating calls (*i.e.* by the logical rules of the model) and by **updating modes** which define the synchronisation between concurrent updating calls. Various types of updating modes have been introduced, with most software tools focussing on a specific subset. BioLQM aims to provide an extensive set of updating modes in a single toolkit. In the following subsections, we further distinguish deterministic and non-deterministic simulations and provide an overview of all updating modes implemented in bioLQM. While stable states, which have no transition toward other states, do not depend on the updating mode, reachability properties and cyclical attractors can be strongly affected by the choice of updating mode as illustrated in Figure 2. More formal definitions of the updating calls and updating modes are given in appendix B.

**Figure 2:**
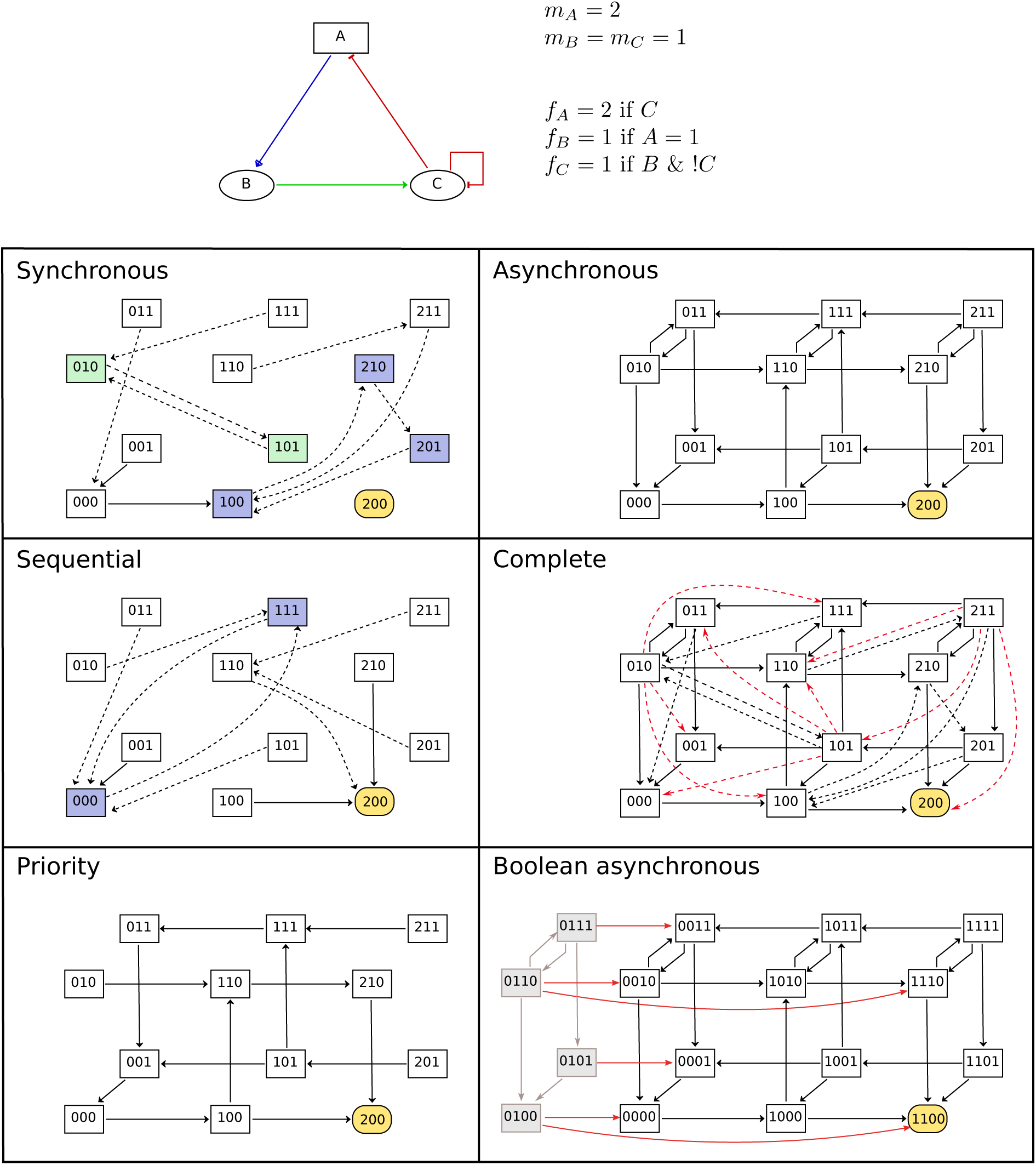
Comparison of updating modes. State transition graphs obtained with the multi-valued model shown in the top part using various deterministic (left-side panels) and non-deterministic (right-side panels) updatings. Dashed arcs denote multiple transitions and node colouring emphasises attractors. Note that the stable state is common to all updatings. The sequential and priority updaters follow the implicit ordering of the components. The STG obtained with the complete updating contains all synchronous and asynchronous transitions, as well as additional transitions to leave the states encompassing more than two updating calls. These transitions are coloured in red in the corresponding panel. Finally, the bottom-right panel contains the asynchronous STG obtained for the booleanized version of the model. In this STG, grey nodes and arcs on the left side correspond to non-admissible states and transitions between them. These states are unreachable from the admissible ones, and the transitions enabling to leave this set of states are highlighted.

### Deterministic simulations

In a **deterministic** simulation, each state has a unique successor, except stable states which have no successor at all as we consider here that a successor must denote a change of state. Starting with an initial state, a deterministic simulation yields an ordered list of successive states, called a **trace**. Given a sufficient number of steps, all traces end in an **attractor**, which can be either a stable state or a **cyclical attractor** of length *k* in which the *k*-th successor of each state is itself. The `trace` tool, illustrated in appendix D, uses an initial state and a deterministic updater to compute a simulation trace. The following deterministic updaters are supported:

- In the classical **synchronous** updating (also called parallel), all logical rules are applied at the same time [16].
- A **sequential** updating applies all rules in a pre-determined order. Instead of evaluating all rules on the original state before updating all components at once as in the synchronous case, they are evaluated on the state obtained after applying the previous rule. The selected order can then change dramatically the successor state: a different sequential updater can be defined for each possible ordering.
- The **block-sequential** updaters generalizes the sequential updaters by considering groups of components updated synchronously [29]. The definition of this updater relies on an ordered partition of the model components.
- The **synchronous priority** updating is also based on a partition of components into blocks, but only the first block containing updated components will be considered. These updatings are a subset of the priority-based updatings introduced by Fauré et al. [11].

### Non-deterministic simulations

In a **non-deterministic** simulation, each state can have several successors. Starting with an initial state, a non-deterministic simulation can lead to a large number of alternative trajectories. This type of dynamics if often represented as a **State Transition Graph** (STG), where the nodes are states of the model, and arcs denote possible transitions between these states. Like in the deterministic case, all trajectories end in an attractor, but starting from an initial state, a non-deterministic simulation can lead to several alternative attractors. These attractors can be **stable states**, **cyclical attractors**, as well as sets of intertwined cycles called **complex attractors**. More formally, all attractors are terminal strongly connected components of the STG. State Transition graphs can represent deterministic traces as well as more complex dynamical behaviours. Such a graph can cover several alternative initial states or even all possible states. The current version of bioLQM supports the definition of non-deterministic updaters, enabling the computation of the lists of successor states. However, it does not provide a complete engine for non-deterministic simulations, or a data structure for state transition graphs. GINsim [25] implements these features on top of bioLQM. The following non-deterministic updaters are supported:

- In the **asynchronous** updating, all logical rules are applied independently. All successors of a state differ from it on exactly one component [37].
- In the **complete** updating, any number of components can be updated at once. This set of successors includes all asynchronous successors, as well as the synchronous one.
- The **priority** updater generalizes the *synchronous priority* introduced above by allowing some of the partitions (priority classes) to be updated asynchronously [11].

### Stochastic simulations

Stochastic updaters enable the computation of a single succesor, which is selected randomly among multiple possibilities and can thus change between calls. A stochastic updater can be derived from any non-deterministic updater by assigning identical probabilities to all transitions defined by the original updater. Alternatively, a custom updater can be constructed by defining individual probabilities.

BioLQM provides the `random` tool to compute single random trajectories using the above stochastic updaters.

This tool is limited to the construction of individual trajectories and does not provide a complete stochastic analysis. As listed in Table 1, bioLQM enables the conversion of Boolean models to the format of the MaBoSS software, which uses the Gillespie algorithm to estimate the probabilities of Boolean states of a continuous time Markov process, and provides a collection of scripts to further analyse the simulation results [35].

### Identification of attractors

The dynamical analysis of large regulatory networks through model simulation suffers from combinatorial explosion, especially in the non-deterministic case. BioLQM implements two published methods based on constraint-solving for the identification of attractors without explicit state enumeration.

1. The first method enables the identification of **stable states** (fixed points) by extracting and combining stability conditions from the logical rules [21]. BioLQM includes this implementation, using decision diagrams to manipulate stability conditions, and introduces an alternative implementation based on the clingo ASP solver [13], which tends to be slower for small models, but can scale better in some cases. Similar methods are also available in the GNA and Pint tools [3, 28].
2. The efficient identification of cyclical attractors and complex attractors remain a challenging problem, especially as these attractors can depend on the updating mode. **Stable patterns** have recently been proposed as an approximation of complex attractors, which can be identified efficiently and does not depend on the updating mode [17, 40]. Here, a pattern is a partially-defined state where some components have a fixed activity level, while others are undefined. Such a pattern represents all states with matching activity levels for the defined components (*i.e.* 2^*k*^ possible states for *k* undefined Boolean components). A pattern is stable if the images of all included states belong to the pattern (the image of a state is its successor in a synchronous updating). BioLQM proposes an adapted version of the method implemented in PyBoolNet [17, 18] using the clingo ASP solver [13], and introduces a new alternative implementation based on decision diagrams.

While complex attractors are well estimated through stable patterns, their exact identification requires further analysis using external software tools, adapted to the selected updating mode. In the synchronous case, the BNS tool [10] uses a SAT solver to identify cyclical attractors of length *k*. This approach could be extended to other deterministic updatings, but can not handle non-deterministic cases. In contrast, BoolSim uses symbolic exploration for the identification of complex attractors in the synchronous and asynchronous case [12]. While this approach scales better than simple simulation, it is more sensitive to combinatorial explosion than approaches based on constraint-solving. To perform the analysis provided by the BoolSim and BNS tools, bioLQM can convert models to their respective formats.

## 4 Model modifications

Several software tools propose to emulate biological **mutations** by generating a model variant in which one or several logical rules have been modified. In bioLQM, such perturbations are a special case of **model modifications**, which consist in computing a new, “modified”, model based on an input model and some parameters if needed. While a perturbed model has the same set of components than the original one, other model modifications can have a different set of components. Each modifier can be described by a keyword (identifier for the type of modifier) and an optional string describing the parameters. Using the model modifier API, bioLQM allows to chain a variety of these modifications before model conversion or analysis. The following describes the various types of model modifications implemented in bioLQM.

### Perturbations

A **perturbation** (often called **mutation**) enables to change some of the logical rules of a model. BioLQM provides three types of “atomic perturbations” (fixed value, range restriction, and removal of a regulator) which modify a single logical rule, while “multiple perturbations” combine several atomic perturbations. Atomic perturbations are briefly described below, more formal definitions can be found in appendix C. The definition of these perturbations is supported by a simple syntax, as illustrated in appendix D and described in the online documentation.

Perturbations are commonly used to model gene knockouts by **blocking the activity level** of the corresponding component to 0, or ectopic expressions by blocking it to 1. Multi-valued components can also be blocked to a higher activity level (inside their normal acticity range).

**Restricting the activity level** of multi-valued components to a smaller range enables to account for a partially impaired activity (loss of the higher activity level(s)) or to set a minimal activity level.

Lastly, it is possible to define the **perturbation of a single interaction**, *i.e.* to remove one of the regulators of a component. This type of perturbation enables for example the definition of the loss of a single binding site preventing the action of the source component on a subset of its targets. The removal of an interaction amounts to rewrite the logical rule of its target component. Note that the atomic perturbation describes the effect on a single target: a single “biological mutation” may correspond to a “multiple perturbation” in the model if several targets are affected by the loss of the same binding site. This type of perturbation is also convenient to evaluate the importance of an interaction representing an hypothetical effect.

### Model reduction

Model reduction aims to ease the analysis of models with a large number of components by constructing a smaller model involving fewer components, but exhibiting similar dynamical properties. BioLQM provides a model reduction method which updates the logical rules of the remaining components to emulate the effect of the removed components [24, 39]. This reduction preserves key dynamical properties of the model, in particular the stable states and stable patterns. However, it can affect some dynamical properties, depending on the choice of reduced components.

This modifier usually relies on the specification of the set of components to reduce. Some types of reduction can be fully automated. In particular, bioLQM supports the reduction of output components, which was shown to preserve attractors and reachability properties [20], as well as the propagation of fixed components, which has also been shown to preserve attractors [32].

After reduction, the reduced components are not fully eliminated from bioLQM: they are no longer allowed to regulate other components, but they keep a logical rule to allow the computation of their expected value in the reduced model.

### Boolean mapping of multi-valued models

As discussed above, some software tools and formats are limited to Boolean models, for example as they rely on specific theoretical results or data structures. To apply such software tools to the analysis of a multi-valued model, we can construct a Boolean model such that its dynamical properties can be transfered to the original multi-valued model.

This **model Booleanization** step is based on the Boolean mapping discussed by Didier, Remy, and Chaouiya [9]. In this mapping, a multi-valued component with a maximal activity level *m* is replaced by *m* Boolean components, each denoting increasing activity. All possible states of the original model can then be associated to states of the Boolean model. The logical rules of the new model ensure that we obtain the same transitions between these states. However, some states of the Boolean model are not mapped to states of the original model. These additional states are called “non-admissible states”.

The dynamical properties observed on the admissible states of the Boolean model can be transfered to the original model. The implementation proposed here further ensures that all synchronous and asynchronous trajectories starting in with a non-admissible state end in an admissible state after a sufficient number of steps. This property ensures that no attractor contains any non-admissible state (see Figure 2).

Model Booleanization is used automatically when converting multi-valued models to formats supporting only Boolean models. It can also be performed explicitely, like other model modifications.

## 5 Usage

BioLQM is freely available under the LGPL v3 license. Documentation, source code and releases are available on http://colomoto.org/biolqm. It is implemented in the Java programming language and thus requires a Java Runtime Environment (JRE 8). A Java compiler (JDK 8) and the Maven build tool are needed to build it from source.

BioLQM is distributed as a JAR file, which can be launched with the command `java-jar bioLQM.jar`. It is also part of the `conda`^2^ package for `GINsim` which was created for the CoLoMoTo Docker image [26]. This package further includes a `bioLQM` command for ease of use.

Hardware requirements strongly depend on the size and structure of models and the operations performed. The complexity of individual logical rules can be a limiting factor: components with tens of regulators could have intricate rules that scale badly. Fortunately, such rules are seldom used in biological models. Any desktop computer should be able to load and convert most models, including large ones. However, the detailled dynamical analysis of models beyond 30 components can rapidly fill the available memory.

The **command-line interface** provides an easy way to convert a model to a different format or to launch a single analysis. It further supports the definition of one or several model modifiers, allowing to save a modified model or to analyse it on the fly. Some examples of command lines are given in appendix D and in the online documentation.

More complex analysis tasks can use the integrated **scripting interface**. Based on the java scripting engine, it supports scripts written in javascript (as part of the java platform) or in another supported language by providing additional libraries (including python and lua among others). An example of script is proposed in appendix E.

### Summary and discussion

The increasing use of logical models of biological regulatory networks led to the development of multiple complementary software tools for their analysis. The recent introduction of the SBML qual format [7] and the formation of the CoLoMoTo consortium [22] aims to facilitate the exchange of models between tools. The bioLQM toolkit enables the use of additional software tools through conversion to their native formats. It provides model conversion operations in the CoLoMoTo notebook [26], enabling the delineation of analysis workflow involving a series of different tools.

BioLQM can also be used to apply various perturbations to the converted models, enabling the study of model variants emulating a knockout, an ectopic activity, or the loss of an interaction. Model modifications include the booleanization of multi-valued models for analysis with tools restricted to a Boolean formalism, as well as model reduction, decreasing the number of components to ease the analysis of complex models.

Finally, bioLQM provides several internal tools for the dynamical analysis of logical models. The first two tools allow the construction of deterministic and stochastic simulation traces, based on a comprehensive collection of updating modes. BioLQM also implements non-deterministic updating modes, which can be used as core components of complete simulation engines, as done by the GINsim [25] and Epilog [38] software suites. Two other tools enable the efficient identification of stable states and the approximation of most complex attractors.

The features described above are organized in a flexible architecture to facilitate the addition of new modules (file formats, model modifications, analysis tools) and to provide a consistent API. In the next version, the configuration API of analysis tools will be improved to provide a better integration with python scripts for use with the new CoLoMoTo notebook. Future work also includes improvements of the representation of the logical rules, which currently rely on decision diagrams (used by the current analysis tools).

Like most modelling tools, bioLQM is currently centered on logical rules, however a complete model may contain important additional information, such as annotations and graphical layout information. The later can be stored along with SBML qual models by using the SBML layout extension. This extension is currently ignored by bioLQM but supported by JSBML and GINsim, it will be supported by future versions of bioLQM. Model annotations are directly supported in SBML core (without additional extensions), however annotations can be defined in any format, hindering interoperability. Further discussions are needed within the community to delineate best practices and ensure that annotations can be shared efficiently.

The reproductibility of model analysis relies on sharing both the model itself and the definition of simulation parameters, in particular initial states and updating modes. A single initial state can be defined in the SBML qual file. Additional initial states and simulation parameters fall in the scope of the Simulation Experiment Description Markup Language (SED-ML) format [4], which does not yet support qualitative models. Ongoing discussions should lead to extensions of the SED-ML format and the Kinetic Simulation Algorithm Ontology [8] to describe model modifications and simulation parameters. These extensions will then be integrated into bioLQM and other qualitative modelling software.

## Author Contributions

AN designed and implemented the software and wrote the paper.

## Acknowledgments

This work benefited from the feedback of members of the CoLoMoTo consortium (colomoto.org). Pedro Monteiro, Claudine Chaouiya and Denis Thieffry provided feedback for the integration in GINsim and Epilog. Julien Dorier and Gautier Stoll helped with the BoolSim and MaBoSS formats respectively. Loic Paulevé provided feedback and implemented the Pint format. Francisco Plana implemented the block-sequential updater. Hannes Klarner implemented the CNET and bnet formats. Celine Hernandez tested the API and provided feedback.

## A Logical Qualitative Model

A **Logical Qualitative Model 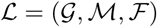** with *n* components is defined by:

- 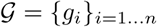, the set of its components.
- 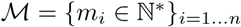, the maximal activity levels of these components, 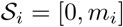 is the activity range of the component *g*_*i*_, and 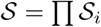 is the state space.
- 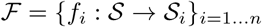, the logical functions defining its dynamical behavior.

A **Boolean model** 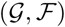 is a logical qualitative model such that all *m*_*i*_ = 1 *∀i ∈* [1, *n*] 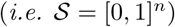.

## B State space and dynamics

In the following, we define the state and dynamics of a model, which can be computed from its logical functions *f*_*i*_. For simplicity, we use *i* as a shorthand for *g*_*i*_ when no ambiguity is possible.

A (qualitative) state 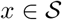 of the model is a vector giving the activity levels of all components, where *x*_*i*_ denotes the activity of the component 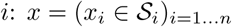. A component *i* is **called to update** in the state *x* if *f*_*i*_(*x*) ≠ *x*_*i*_. The state *x* is a **stable state** (also called fixed point, or steady state) if no component is called to update, *i.e.* if *x*_*i*_ = *f*_*i*_(*x*) ∀*i ∈* 1 *… n*.

We define unitary update functions 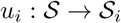 such that *u*_*i*_(*x*) = *x*_*i*_ + ∆_*i*_(*x*), where

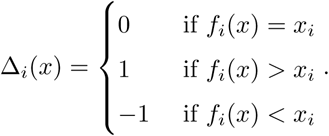

In the Boolean case, these unitary functions are identical to the logical functions *f*_*i*_ by construction. They enforce unitary transitions in the multi-valued case.

We call *f* (*x*) = (*f*_*i*_(*x*))_*i*=1*…n*_ the **image** of the state *x*, and *u*(*x*) = (*u*_*i*_(*x*))_*i*=1*…n*_ its **unitary image**.

Given a state 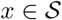 and a subset of the components 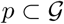, let 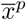 be the state where all the components of the subset have been updated at once:

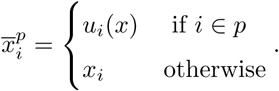

For simplicity, 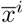 denotes 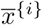.

### Deterministic updatings

- In the **synchronous** updating, the unique successor of *x* is its unitary image *u*(*x*).
- In the **sequential** updating following the default order of components, the unique successor of the state *x* is (*u_n_ ◦ ⋅ ⋅ ⋅ ◦ u*_1_)(*x*). A different sequential updater can be defined for each possible ordering of 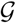.
- A **block-sequential** updater is based on an ordered partition of 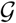 in *m* non-overlapping subsets (*m ≤ n*). Let 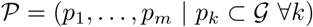 be this ordered partition. The function *v*_*k*_ : 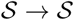 such that 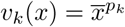 updates all components of the subset *p*_*k*_ synchronously. The unique successor of the state *x* by the block-sequential updater defined by the ordered partition 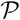 is then given by combining these functions: (*v_m_ … ◦ v*_1_)(*x*).
- The **synchronous priority** updating is based on the partitioning 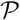 defined for the block-sequential case, but only the first updated subset is considered. If the state *x* is not a stable state, then there exists *k* such that the subset *p*_*k*_ is the first one containing updated components: *v*_*k*_(*x*) ≠ *x* and *v*_*i*_(*x*) = *x* ∀*i* < *k*. Then *v*_*k*_(*x*) is the successor of *x* in this updating. Note that this type of updating is the deterministic subset of the non-deterministic priority updatings defined below.

### Non-deterministic updatings

1. In the **asynchronous** updating, all logical functions are applied independently and all successors of the state *x* differ from *x* by exactly one component: the set of successors of *x* is 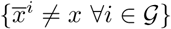.
2. In the **complete** updating, any number of components can be updated at once: the set of successors of *x* is 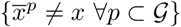. This set of successors includes all asynchronous successors, as well as the synchronous one.
3. The **priority** updater generalizes the *synchronous priority* defined above by considering asynchronous updates between the blocks (priority classes). Starting with 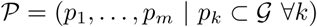 the ordered partition of 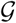 defined above, each subset *p*_*k*_ is further partitioned in *a*_*k*_ subsets *p_k,_*_1_*, …, p_k,ak_*. If the state *x* is not a stable state, then there exists *k* such that the subset *p*_*k*_ is the first one containing updated components: *v*_*k*_(*x*) ≠ *x* and *v*_*i*_(*x*) = *x ∀i < k*. Then *v_k,_*_1_(*x*)*, …, v_k,ak_*(*x*) are the *a*_*k*_ successors of *x* in this updating.

## C Model modifications

### Range restriction

We call 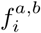 the restriction of the function *f*_*i*_ to the range [*a, b*] (with 0 *≤ a ≤ b ≤ m_i_*) such that:

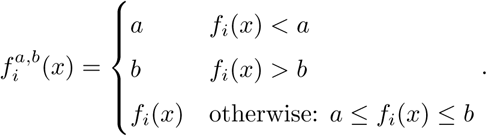

### Perturbation of an interaction

The removal of an interaction (*i, j*) (*i.e.* the removal of *i* from the regulators of *j*) further requires the specification of 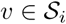 and leads to the modification of *f*_*j*_, the logical rule associated to *j*.

Let 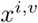 be the state such that 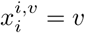 and 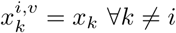. The perturbation constructs the modified function 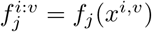.

## D Command-line examples

Launching bioLQM without arguments displays a help message, listing all available features:

~~~
bioLQM
~~~

Converting a model in the SBML qual format to the BoolSim format:

~~~
bioLQM model.sbml model.boolsim
~~~

Loading a model, applying a knockout (of the `Akt` component) and saving the resulting modified model:

~~~
bioLQM model.sbml -m perturbation:Akt%0 perturbed.sbml
~~~

The syntax for model perturbations is further described in the online documentation. The modified model can also be saved to a different format (combining model modification and conversion):

~~~
bioLQM model.sbml -m perturbation:Akt%0 perturbed.boolsim
~~~

Loading a model and computing the stable states:

~~~
bioLQM model.sbml -r fixpoints
~~~

In this more complex example, the loaded model is perturbed before being reduced, then the stable states of the resulting model are computed:

~~~
bioLQM model.sbml -m perturbation:Akt%0 -m reduce:fixed -r fixpoints
~~~

## E Scripting example

Calling bioLQM in script mode:

~~~
bioLQM -s generate_perturbations.js model
~~~

Content of the script file for the generation of all possible knockout perturbations:

~~~
filename = lqm.args[0]
model = lqm.load(filename)
nodes = model.getComponents()
for (i in nodes) {
    node = nodes[i]
    perturbed = lqm.modify(model, ‘perturbation’, node+’%0’)
    lqm.save(perturbed, filename+"_"+node+"_KO.sbml")
}
~~~

## Footnotes

1 See http://colomoto.org/biolqm/

2 https://conda.io

